# Extraction of ultrashort DNA molecules from herbarium specimens

**DOI:** 10.1101/076299

**Authors:** Rafal M. Gutaker, Ella Reiter, Anja Furtwängler, Verena J. Schuenemann, Hernán A. Burbano

## Abstract

DNA extracted from herbarium specimens is highly fragmented and decays at a faster rate than DNA from ancient bones. Therefore, it is crucial to utilize extraction protocols that retrieve short DNA molecules. Improvements in extraction and library preparation protocols for animal remains have allowed efficient retrieval of molecules shorter than 50 bp. We adapted those improvements to extraction protocols for herbarium specimens and evaluated their performance by shotgun sequencing, which allows an accurate estimation of the distribution of fragment lengths. Extraction with PTB buffer decreased median fragment length by 35% when compared to CTAB. Modifying the binding conditions of DNA to silica allowed for an additional decrease of 10%. We did not observe a further decrease in length when we used single-stranded instead of double-stranded library preparation methods. Our protocol enables the retrieval of ultrashort molecules from herbarium specimens and will help to unlock the genetic information stored in herbaria.

## Method summary

We optimized the extraction procedure for isolating ultrashort DNA fragments from herbarium specimens through combination of PTB lysis buffer and modifications previously used for ancient bones. We show the advantage of this protocol over others by estimating the DNA fragment length through shotgun sequencing.

Since ancient DNA (aDNA) is highly fragmented, it is particularly important to employ extraction protocols that retrieve ultrashort molecules (< 50 bp). It has been shown that a recently developed extraction protocol for animal remains efficiently recovers those molecules (1, which has allowed sequencing highly fragmented hominin (2) and cave bear remains (1) that are hundreds of thousands of years old. DNA retrieved from herbarium specimens is also highly fragmented because it decays six times faster than in bones (3). Consequently, DNA from century-old herbarium specimens is as short as that of thousands of years old animal remains. To take full advantage of the genetic information stored in those samples it is important to optimize the extraction of ultrashort molecules from desiccated plant tissue.

We assessed the impact of extraction and library preparation methods on the distribution of DNA fragment lengths in 20 *Arabidopsis thaliana* herbarium specimens, which were collected between 1839 and 1898 (Table S1). We used a hierarchical experimental design that includes three different phases due to limited availability of tissue per sample (Figure 1). In phase one and two we used 10 *A. thaliana* samples (~20 mg of leaf tissue each), which were subjected to two different extraction protocols (~10 mg of tissue per treatment), followed by double-stranded library preparation. To compare the performance of double-and single-stranded library preparation methods, in phase three we applied single-stranded library preparation method to DNA extracts produced by the most efficient DNA extraction protocol in phase two. In each phase we evaluated the performance of the methods by sequencing the genomic libraries with the Illumina MiSeq platform (Table S2).

**Figure 1.**
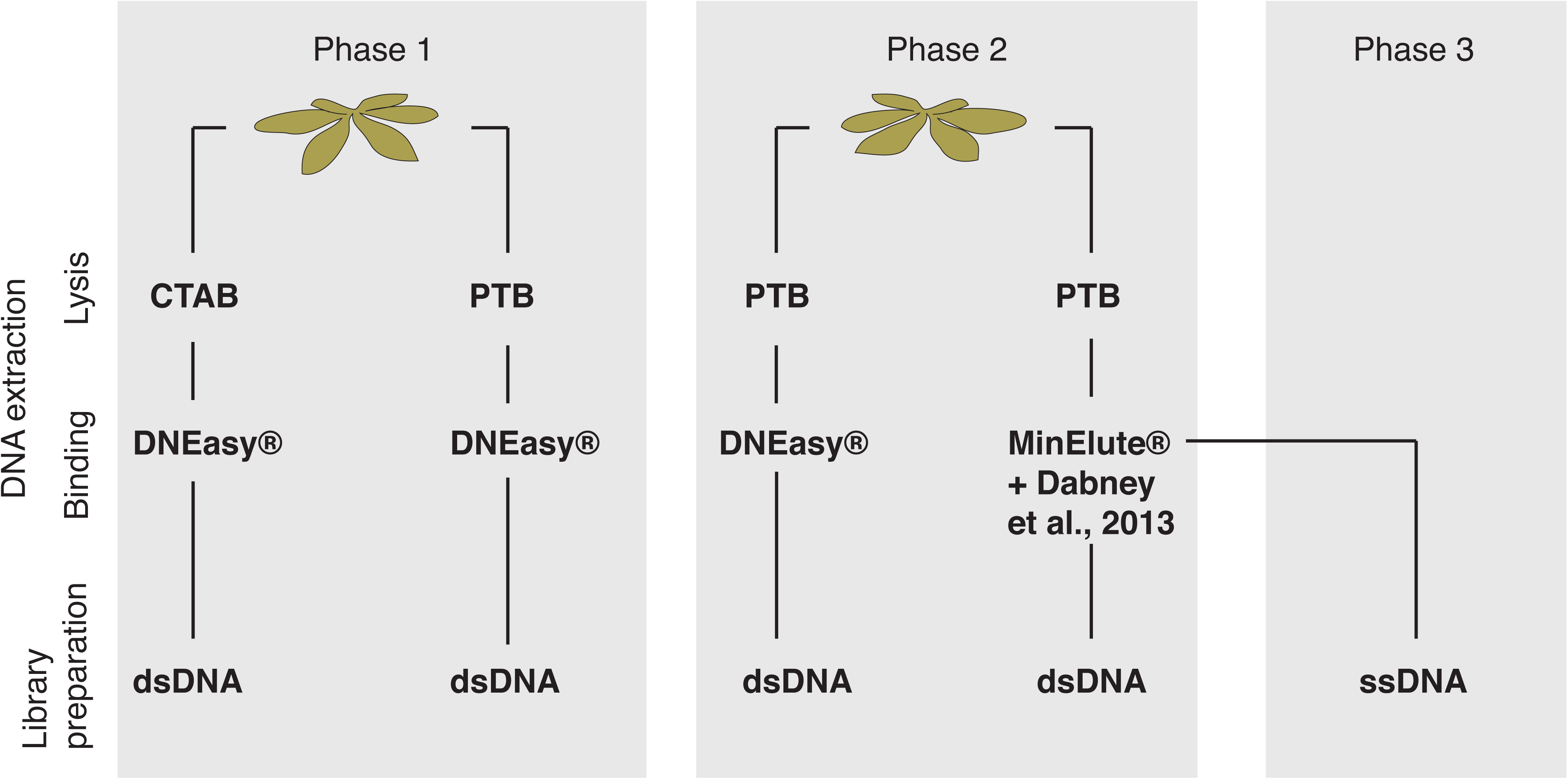
Experimental design for testing the effect of DNA extraction and library preparation protocols on properties of sequenced libraries from herbarium specimens. Experiments were conducted in three phases. In phase one we subject 10 herbarium specimens of *Arabidopsis thaliana* to extraction with two different lysis buffers and compare sequencing results. In phase two we tested two DNA-binding methods on second set of 10 *A. thaliana* specimens. In phase three we compared the libraries constructed with double-and single-stranded methods.

Extraction buffers used for ancient bones and teeth are commonly composed predominantly or exclusively of EDTA and proteinase K (4), reagents that are not optimal for DNA extraction from plant tissue. Hence, in the first phase we tested two commonly used DNA extraction buffers for historical plant specimens, which contain either cetyl-trimethyl ammonium bromide (CTAB), or a mixture of N-phenacylthiazolium bromide (PTB) and dithiothreitol (DTT) (5) (Figure 1). CTAB is a strong detergent that under high salt concentrations binds to polysaccharides and aids their removal from the solution (6). Although CTAB is highly used in DNA extractions from modern plants, it has been shown that it does not have a detectable effect when applied to non-carbonized archaeobotanical remains (7). PTB is a substance that cleaves glucose-derived protein cross-links (8) and can help to release DNA trapped within sugar-derived condensation products (9); it has been effectively used to retrieve DNA from archaeobotanical remains (10). DTT digests disulfide bonds releasing thiolated DNA from cross-link complexes (11). In order to allow better comparison of the CTAB and PTB protocols, we replaced the ethanol precipitation step of the CTAB method with silica column binding (12) provided with the DNeasy^®^ Plant Mini Kit. Subsequently, libraries were prepared using a double stranded DNA library protocol (13).

Based on qPCR measurements on unamplified libraries, the PTB protocol recover a higher number of unique library molecules than CTAB protocol (paired t-test p = 0.007) (Figure 2F and Table S3). We found that PTB decreases the median fragment length by 35% (from 88 to 57 bp) (paired t-test p = 2.8e-06) when compared to CTAB (Figure 2A and 2B). This decrease in length was also manifested as a higher proportion of damaged sites (lambda) (paired t-test p = 1.3e-06) (Figure 2C), which represents the fraction of bonds broken in the DNA backbone (14, 15). In addition, DNA molecules extracted with PTB buffer showed more cytosine (C) to thymine (T) substitutions at the 5’ end (paired t-test p = 1.2e-06; Figure 2G). C-to-T substitutions are typical damage patterns of aDNA and result from spontaneous deamination of C to uracil (U), which is read as T by the polymerase (16, 17). It is possible that shorter and more damaged fragments of DNA were released after cross-links were resolved by PTB and DTT, since there is a strong negative correlation between median fragment length and C-to-T substitutions at first base (R^2^ = 0.44; p = 1.5e-07; N = 50) (Figure S4). Alternatively, the observed variation in fragment length distribution could be explained by unknown chemical incompatibilities of lysis and binding buffer, i.e. certain reagents could in principle reduce DNA-binding properties of the buffer. Finally, in the CTAB protocol we apply a chloroform-isoamyl alcohol wash, which could also reduce recovery of short molecules.

**Figure 2.**
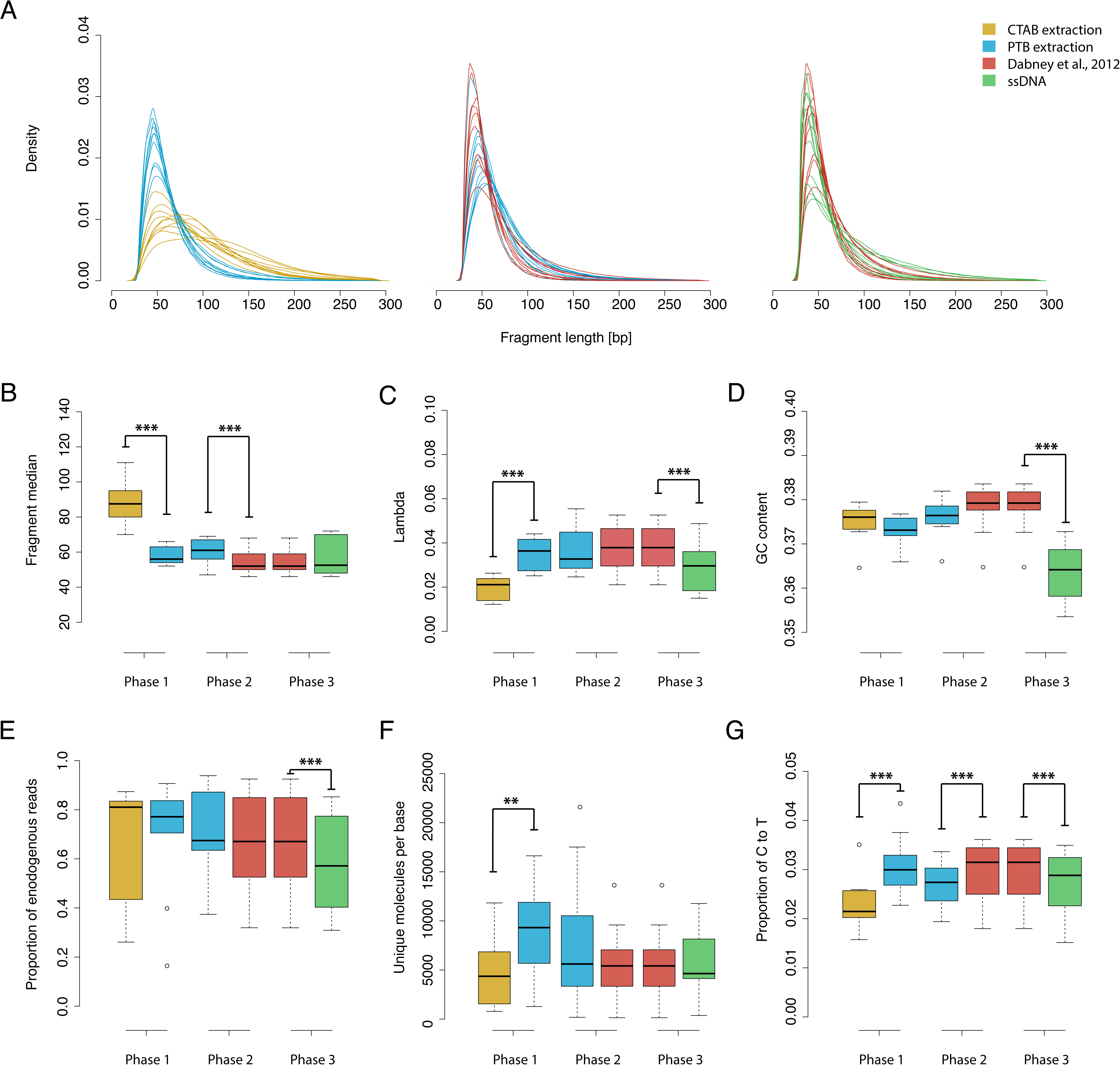
The effect of DNA extraction and library preparation protocols on different properties of DNA sequencing libraries. The figure depicts the results from experiments in phases 1-3 (Figure 1). (A) Distribution of fragment lengths of merged reads mapped to the *Arabidopsis thaliana* reference genome. The y-axis shows the kernel density estimates. (B-G) Distributions represented as box and whisker plots; medians are depicted by thick black lines, boxes represent data between quartile Q1 and Q3, whiskers extend to 1.5 times the interquartile range between Q1 and Q3, and points symbolize outliers. Comparisons within experiments that result in significant differences in a paired t-test are connected with black lines (‘***’ indicates an alpha level of 0.005) (B) Fragment length medians. (C) Proportion of broken DNA fragments (lambda). (D) Proportion of GC content. (E) Proportion of endogenous DNA (proportion of reads mapped to *A. thaliana* reference genome). (F) Number of unique molecules per base of *A. thaliana* reference genome (molecule coverage of DNA extract) calculated from qPCR measurements on unamplified libraries. (G) Percentage of cytosine to thymine substitutions at first base at the 5’ end.

In the second phase, to further increase the recovery of short fragments, we used PTB/DTT, which was the most successful extraction buffer in phase 1, and evaluated two systems for binding DNA to silica. We tested DNeasy® mini spin columns (Qiagen) in combination with the binding buffer used in the Plant Mini kit and MinElute® silica spin columns in conjunction with a binding buffer optimized for the recovery of short molecules from animal remains (1) (Figure 1). We found that the latter method decreased the median fragment length by 10% (from 60 to 54 bp) (paired t-test p = 1.9e-04), which shows that it is suitable to recover very short sequences also from herbarium specimens (Figure 2A and B). The frequency of C-to-T substitutions at the first base differed significantly between the two DNA binding methods (paired t-test p = 3.3e-03) (Figure 2G), with a decrease in median fragment length again being accompanied by an increase in C-to-T substitutions.

To investigate whether library preparation has an effect on fragment length distribution, in the third phase we produced single-stranded DNA (ssDNA) libraries using the extracts from the modified PTB/DTT extraction (18,19) and compared them to the dsDNA libraries constructed from the modified PTB extraction material of phase 2 (Figure 1). We did not observe a significant decrease of the median of the fragment length distribution in ssDNA libraries (paired t-test p = 0.44) (Figure 2A and B). Instead, the shape of the distribution changed towards larger numbers of longer and shorter molecules at the cost of intermediate-size molecules, which is reflected in decreased lambda (Figure 2C) and congruent with previous findings (19). Similarly to Gansauge and Meyer (2013), we also detected a reduction of GC content in ssDNA libraries when compared to dsDNA (Figure 2D). This phenomenon can be attributed to a known bias in dsDNA libraries towards molecules with higher GC content (20,21). We detected uniform GC content across the distribution of fragment lengths, which suggests that the ssDNA library preparation protocol excels in reducing those biases (Figure S8). In contrast to previous reports (19), the ssDNA library method did not produce an increase in the proportion of endogenous DNA (Figure 2E, Figure S1, S6). However, it has been suggested that increase in the proportion of endogenous DNA occurs only when the initial content of endogenous DNA is lower than 10% (22, 23). Our *A. thaliana* samples have endogenous DNA between 16% and 94%, which could explain why we did not detect a gain in endogenous DNA.

In summary, we demonstrate that the choice of extraction buffer has a great impact on the length distribution of molecules recovered from herbarium specimens. Ultrashort molecules are most efficiently retrieved using a combination of PTB/DTT mixture for DNA extraction and the buffers and conditions suggested by Dabney *et al*. (2013) for DNA binding. The two library preparation methods tested here appear to be equally efficient in retaining short DNA fragments, however, while single stranded method reduces GC bias in library it also decreases the fraction of endogenous DNA. We present the DNA extraction protocol that increases the recovery of short fragments and thus the accessibility of precious herbarium specimens for genetic analyses.

## Author contributions

R.M.G, V.J.S, and H.A.B designed the experiments. R.M.G., E.R. and A.F. performed the experiments. R.M.G analyzed the data. R.M.G and H.A.B wrote the manuscript with contributions from all authors.

## Acknowledgements

We thank curators from the Missouri Botanical Garden, University of Illinois, National Museum of Natural History, West Virginia University, Harvard University and New York Botanical Gardens for kindly providing samples for this study; Marco Thines, Charles B. Fenster and Matthew T. Rutter for sampling the herbarium specimens; Daniel Koenig for advice in the experimental design; Patricia Lang, members of the Research Group for Ancient Genomics and Evolutions, and especially Matthias Meyer for comments on the manuscript. This work was funded by the Presidential Innovation Fund of the Max Planck Society.

